# Measuring significant changes in chromatin conformation with ACCOST

**DOI:** 10.1101/727768

**Authors:** Kate B. Cook, Karine Le Roch, Jean Philippe Vert, William Stafford Noble

## Abstract

Chromatin conformation assays such as Hi-C cannot directly measure differences in 3D architecture between cell types or cell states. For this purpose, two or more Hi-C experiments must be carried out, but direct comparison of the resulting Hi-C matrices is confounded by several features of Hi-C data. Most notably, the genomic distance effect, whereby contacts between pairs of genomic loci that are proximal along the chromosome exhibit many more Hi-C contacts that distal pairs of loci, dominates every Hi-C matrix. Furthermore, the form that this distance effect takes often varies between different Hi-C experiments, even between replicate experiments. Thus, a statistical confidence measure designed to identify differential Hi-C contacts must accurately account for the genomic distance effect or risk being misled by large-scale but artifactual differences. ACCOST (Altered Chromatin Conformation STatistics) accomplishes this goal by extending the statistical model employed by DEseq, re-purposing the “size factors,” which were originally developed to account for differences in read depth between samples, to instead model the genomic distance effect. We show via analysis of simulated and real data that ACCOST provides unbiased statistical confidence estimates that compare favorably with competing methods such as diffHiC, FIND, and HiCcompare. ACCOST is freely available with an Apache license at https://bitbucket.org/noblelab/accost.

## 1 Introduction

An increasing number of experimental techniques—including Chi A-PET [9], Hi-C [13], Hi-ChIP [15], PLAC-seq [7], SPRITE [18], and GAM [2]—allow for the high-throughput characterization of pairwise chromatin contacts. These techniques have helped to elucidate the roles that chromatin 3D architecture play in critical cellular processes such as gene regulation, DNA replication, and splicing. Notably, aspects of chromatin structure have been implicated in the etiology of several disease phenotypes, in which genetic modifications induce changes in chromatin structure, which in turn induce changes in gene expression [12].

Accordingly, a key statistical challenge for 3D chromatin analyses is to assign statistical confidence measures to observed differences in chromatin structure. Such differences may arise between different developmental stages, cell types, individuals or disease states. Empirically, chromatin structure exhibits differences at multiple scales, including differences in nuclear shape and volume, full chromosomes, compartments, topologically associating domains (TADs), sub-TADs, or promoter-enhancer loops. In this work, we focus on calling differential interactions at the finest level, i.e., for a given contact matrix defined with respect to genomic loci (“bins”) of size *w* bp, we ask whether the observed contact count associated with two w-bp loci *x* and *y* in experimental condition *A* is significantly different from the corresponding *(x, y)* contact count from condition *B.*

A successful method for assigning confidence estimates to differential interactions should exhibit four key properties. First, the method must control the false discovery rate (FDR) associated with an accepted set of differential interactions. Controlling FDR, as opposed to family-wise error rate, makes sense in the context of most 3D chromatin studies, which are often hypothesis-generating studies aimed to produce a large set of discoveries. Second, a good method must take into account the particular biases associated with the assay (such as Hi-C) that generated the data. As we will see, accomplishing this goal turns out to be very challenging. Third, because of the high cost associated with generating 3D chromatin data—in particular, the cost of sequencing deeply enough to characterize all pairs of loci in the genome—an ideal method will be capable of leveraging structure in the observed data to enable calling significant differential interactions even when only a single pair of experiments has been performed. Fourth, as with essentially any statistical testing procedure, an ideal method should maximize statistical power; i.e., the method should identify as many differential interactions as possible, given the data, while still controlling the FDR.

Note that, although multiple genome-wide 3D chromatin assays have been developed, most existing statistical testing frameworks focus on the first and still most widely used assay, Hi-C. Accordingly, we focus here on Hi-C analysis, though many of the methods described here may also apply directly to data generated using assays such as ChIA-PET, SPRITE, Hi-ChIP, PLAC-seq, or GAM.

Early Hi-C studies generally employed simple fold-change statistics to identify differential contacts [4, 23], but these methods have subsequently been supplanted by a variety of more statistically sophisticated techniques.

- HOMER [10] implements a simple test for differential interactions, based on binomially-distributed counts. HOMER can only consider comparisons between two libraries and cannot take into account replicate experiments.
- ChromoR [21] carries out variance stabilization of Hi-C data using the the Haar-Fisz transform [8], which transforms Poisson distributed data into Gaussian distributed coefficients at multiple scales. The resulting coefficients are subjected to wavelet-based denoising. ChromoR then analyzes the resulting transformed data using either change-point analysis for the purpose of detecting chromosomal aberrations or domain boundaries, or Bayes factor analysis for the detection of differential contacts.
- HiBrowse [17] is a Python tool with a web interface that employs the edgeR RNA-seq package [20] to compare pairs of Hi-C experiments.
- The R package diffHic [14] also uses the edgeR statistical framework but dispenses with the web interface. DiffHic offers a range of functionality, including mapping, quality filtering, binning, normalization, and estimation of statistical confidences for differential contacts. Notably, the package is able to accommodate complex experimental designs, including paired or blocked designs and experiments involving more than two groups.
- FIND [-5] uses a spatial Poisson model to measure differences in chromatin loops, with the aim of relaxing the independence assumption between adjacent loci.
- HiCcompare [22] normalizes the data and then tests for differential interactions on the normalized data directly, using a Z-score comparison. HiCcompare does not leverage replicate experiments.

For all of these methods, a key challenge is accurately accounting for the unique features of Hi-C data. For example, several widely used methods can normalize Hi-C data to account for positional biases that arise due to GC content, mappability, and density of restriction-enzyme sites along the genome [11, 25]. Variants of these normalization routines are integrated into all of the methods outlined above.

A second prominent and potentially problematic feature of Hi-C data is the “genomic distance effect,” in which pairs of loci that are close together along the genome are very likely to be observed in contact simply due to the random polymer looping behavior of DNA. The genomic distance effect manifests itself most obviously in the form of a marked enrichment of high counts near the diagonal in the Hi-C contact matrix.

This work was primarily motivated by the observation that the genomic distance effect can vary markedly between two Hi-C experiments. The result, in a Hi-C matrix showing log fold-changes in contact counts, is an enrichment of apparently differential contacts far from the diagonal (Figure 1A). Notably, these differences remain even after normalizing the data using a method such as iterative correction and eigenvalue decomposition (ICE) [11] (Figure 1B). The source of this kind of large-scale change in genomic distance effect is not always clear. In some settings, such differences may indicate changes in chromatin compaction [24] or differences in enrichment of various stages of the cell cycle [16]. Alternatively, differences in the genomic distance effect may reflect experimental artifacts: we have observed non-trivial differences even between replicate Hi-C runs (Figure 1C). Regardless of the source of these observed differences, we reasoned that any method aiming to identify local changes in chromatin structure should attempt to ignore large-scale differences in the genomic distance effect.

**Figure 1:**
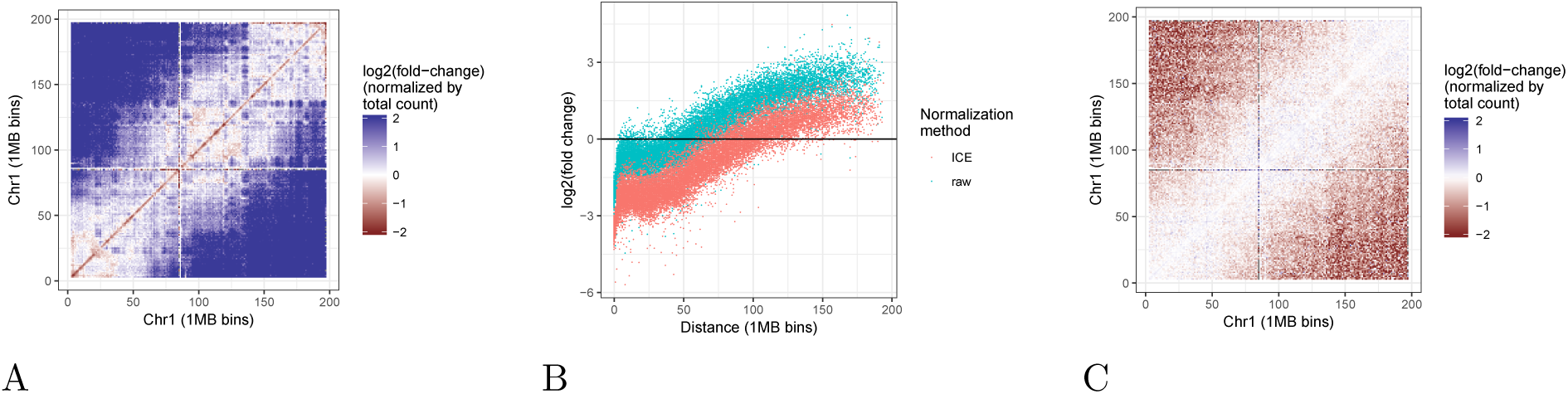
Genomic distance effect leads to artifactual differences in Hi-C contacts. (A) The figure plots the log fold-change in Hi-C contact counts between embryonic stem cell (ESC) and cortex Hi-C data (see Methods for details), plotted at 10 kb resolution on human chromosome 21. (B) Scatter plot of Hi-C log-fold change as a function of genomic distance, where each point corresponds to a pair of 10 kb loci. The two point colors correspond to raw counts (red) and counts normalized using ICE (blue). (C) Similar to panel B, but for two replicate Hi-C experiments performed in mouse cortex.

One other method—HiCcompare, which was published while our method was under development—also attempts to correct for differences in genomic distance effect. HiCcompare works by plotting the log fold change in contact counts as a function of genomic distance, and then fitting a LOESS regression curve to this plot. The regression curve is then used to normalize the observed interaction counts. Differential contacts are then identifying by converting the normalized values to Z-scores and then computing p-values using the standard normal distribution.

In this paper, we introduce ACCOST (Altered Chromatin Conformation STatistics), which assigns statistical confidence estimates to differences in Hi-C contacts, while taking into account differences in the genomic distance effect. Specifically, ACCOST leverages counts at similar genomic distances to estimate the variance of the contact counts, similar to how tools such as DESeq [1] estimate gene expression variance by grouping together genes with similar expression. ACCOST includes distance-dependent normalization factors, thereby allowing for accurate measurement of differential contact counts at short and long ranges. The method accommodates but does not require biological replicates. We demonstrate, via simulation and analysis of real Hi-C data, that ACCOST delivers unbiased statistical confidence estimates while successfully controlling for systematic changes in the genomic distance effect.

## 2 Material and Methods Model

Formally, we consider *m* contact count matrices *C*^*k*^ ∈ ℕ^*n*×*n*^, for *k* = 1,..,,*m*, where 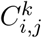 is the interaction count between loci *i* and *j* in the *k-th* experiment (for 1 ≤ *i*,*j* ≤ *n* and 1 ≤ *k* ≤ *m).* Each experiment is done in one of two conditions *𝒜* = {*cond*_1_, *cond*_2_}, and we denote by *ρ*{*k*} ∈ *𝒜* the condition corresponding to the *k-th* contact count matrix, for 1 ≤ *k* ≤ *m.* For each condition *A* ∈ *𝒜*, we denote by *m*_*A*_ = | *ρ*^-1^ (*A*)| the number of contact count matrices done in condition *A;* in particular it holds that Σ _*A ∈ 𝒜*_ *m*_*A*_ *= m.*

We model the count 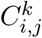 as a negative binomial (NB) random variable with mean 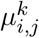 and variance 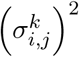. In order to be able to estimate the mean and the variance from few Hi-C replicates, we make the following assumptions, inspired by the assumptions of the DEseq model for RNA-seq data [1]:

- The mean 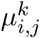 that is the expected number of counts between loci *i* and *j* in the *k-th* matrix corresponding to condition *ρ (k)*, is equal to a quantity <u>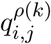</u> which represents the “true” (but unknown) number of interactions between loci *i* and *j* in condition *ρ* (*k*), multiplied by an experiment- and loci-specific factor as follows: 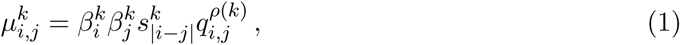

where *β*^*k*^ ∈ ℝ^*n*^ is an experiment-specific vector of loei-speeific biases to account for various biological and technical biases [11] and *s*^*k*^ ∈ ℝ^*n*^ is a vector of distance-specific size factors to account for the experiment-specific “genomic distance effect”. This generalizes the notion of size factors used to correct for library size in RNA-seq [1], but considering each distance and pair of loci separately.
- As in the DEseq model, the variance is the sum of a shot noise term and of a raw variance term:

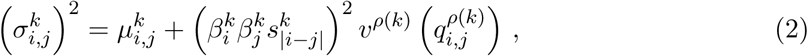

where *v*^*A*^ is a smooth non-negative function, for *A* ∈ *𝒜.*

We now explain how we estimate the parameters *β*, s, *q* and *v* of the model from a set of count matrices *C*^1^,…, *C*^*m*^:

- For the vectors of locus-specific biases *β*^*k*^ (for *k* = 1, …,*m*), we borrow a standard ICE normalization estimator 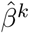 performed independently on each count matrix [11].
- For the distance-specific size factors *s*^*k*^ (for *k* = 1,…, *m*), we enforce that the median normalized count for pairs of bins at each given distance in each matrix is the same, by taking for *d* ∈ [0, *n* − 1] and *k* ∈ [1, *m*]: 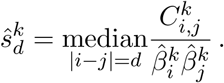
- For the normalized count matrix *q*^*A*^, for *A* ∈ *𝒜*, we first define the normalized count matrices of each experiment: 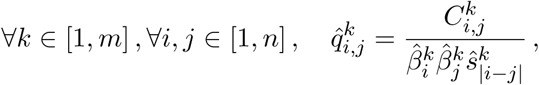

and average these estimates over replicates for each condition: 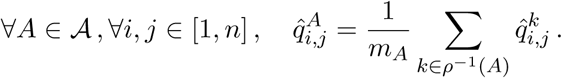
- To estimate the normalized raw variance functions *v*^*A*^ (for *a* ∈ *𝒜)*, we start by estimating the variance of 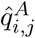 for any *i*,*j* ∈ [1, *n*]. Since we are in situations where the number of replicates *m*,*A* can be small (up to.*m*_*A*_ = 1), we propose to leverage informations within each matrix to estimate this variance. For each bin *(i*,*.j)*, we consider a set of other bins *N*(*i*,*j)* ⊂ [1, *n*]^2^ where we believe 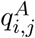 should be almost constant. As an example, we consider *N*(*i*,*j*) to be the set of all other bins (*u, v*) at the same distance, i.e., | *u* − *v* | = | *j* − *i* | or to restrict *N*(*i*,*j*) further by only considering other bins at the same distance but in some window around (*i, j*). Given such a neighborhood, we form the estimates: 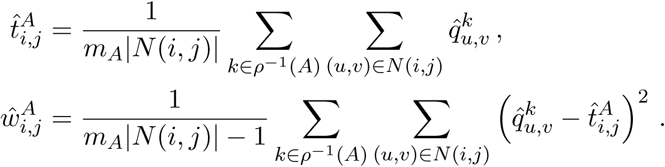 Assuming that 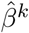 and *ŝ*^*k*^ are exact and nonrandom estimators of *β*^*k*^ and *s*^*k*^, on the one hand, and that 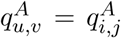 for all (*u*,*v*) ∈ *N*(*i*,*j*), on the other hand, we deduce from our model (1)-(2) that the expectation and variance of 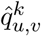 for (*u*,*v*) ∈ *N*(*i*,*j*) and *k* ∈ *ρ*^−1^(*A*) are respectively given by: 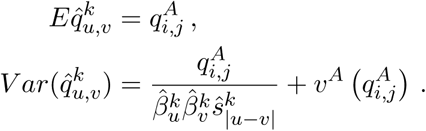 A simple computation shows that if *X*_1_, …, *X*_*N*_ are independent random variables with the same mean and different variances, then: 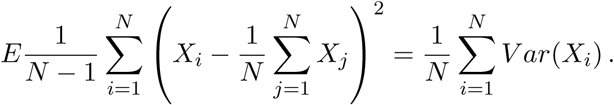 Using this result with the random variables 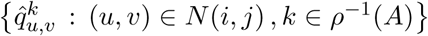 leads to: 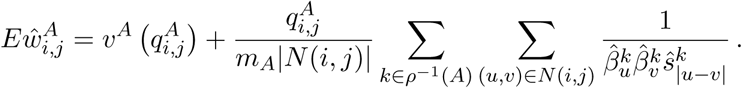 Let us define 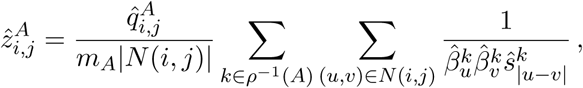

then 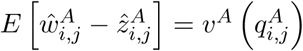, and we can estimate *v*^*A*^ by regressing 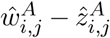 as a polynomial function of 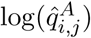, encouraging smoothness. Negative values of *w* − *z* are rare and are filtered out by the algorithm.

To test for differential count between conditions at a given locus (*i, j*), we define as test statistics the total counts in different conditions:

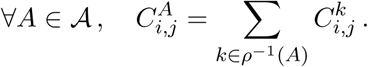

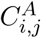 is a sum of *m*_*A*_ independent NB-distributed random variables. We approximate its distribution by an NB distribution with mean and variances:

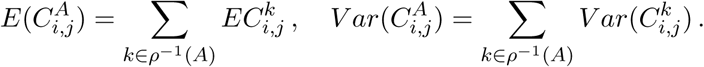

Under the null hypothesis where 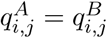, we can get an estimate of the shared normalized count:

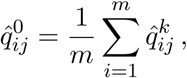

from which we deduce an estimate of the mean and variance of 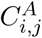 for *A* ∈ 𝒜:

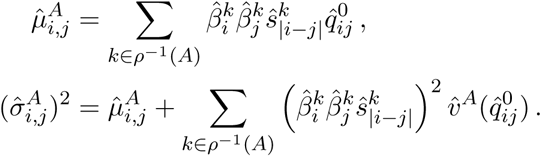

Given two conditions 𝒜 = {*A, B*}, we then follow the technique of DESeq to obtain a p-value by conditioning on the total count 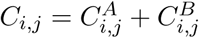:

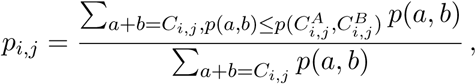

where

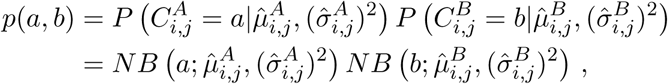

since 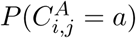 and 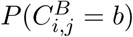 are independent under the null hypothesis.

### 2.1 Data sets

GM12878 and IMR90 Hi-C data at 1Mb and 50kb resolution were downloaded in text form using the `juicer dump` command of Juicer tools [6]. Replicates are numbered as in Table S1 of [19], heading “biological replicate number.”

Processed cardiomyocyte data in count matrix form was obtained from the authors [3].

All datasets were ICE normalized using `iced` (https://github.com/hiclib/iced).

### 2.2 Competing methods

For analysis by diffHiC, contact count matrices were loaded and transformed into ContactMatrix objects. Following the diffHiC documentation, we filtered by average count and applied LOESS normalization before using diffHiC to estimate p-values. As prescribed in the diffHiC manual, we applied LOESS normalization separately to the near-diagonal bin pairs (defined as pairs with genomic distance below 50 Mb) and to other pairs.

Following the FIND documentation, we loaded contact count matrices and ran FIND with parameters windowSize = 3, method = hardCutoff, and alpha = 0.7.

Following the HiCcompare documentation, we loaded contact count matrices and ran HiCcom-pare with parameters A.min = 15 and adjust.dist = true.

## 3 Results

### 3.1 ACCOST correctly accounts for the genomic distance effect

ACCOST extends the statistical model used by DEseq [1] to explicitly capture observed differences in the genomic distance effect between different Hi-C experiments. The key idea is to re-purpose DEseq’s “size factors,” which were originally developed to account for differences in read depth between samples, to instead model the genomic distance effect and the loci-specific biases. The extension is non-trivial because a separate set of factors must be created for each genomic distance (see Methods). To estimate the variance in contact counts in cases where the number of biological replicates is small, we assume that pairs of loci at the same genomic distance act similarly, and calculate, for each genomic distance, sample means and variances for all pairs of loci across all replicates. We then estimate a smooth function of the variance as a function of the mean (Methods) to come to our final estimate of the variance.

Visualization of log-fold change values between two different cell types, mouse ESC and cortex, illustrates the effect of ACCOST normalization. A raw log fold-change comparison (Figure 2A) highlights differences primarily due to the difference in sequencing depth of the two libraries. Cor-recting for this effect by simple linear rescaling leaves a clear pattern associated with the genomic distance effect, with large fold-changes farther from the diagonal (Figure 2B). This pattern is not eliminated by ICE normalization (Figure 2C) but is successfully removed by ACCOST’s normal-ization (Figure 2D).

**Figure 2:**
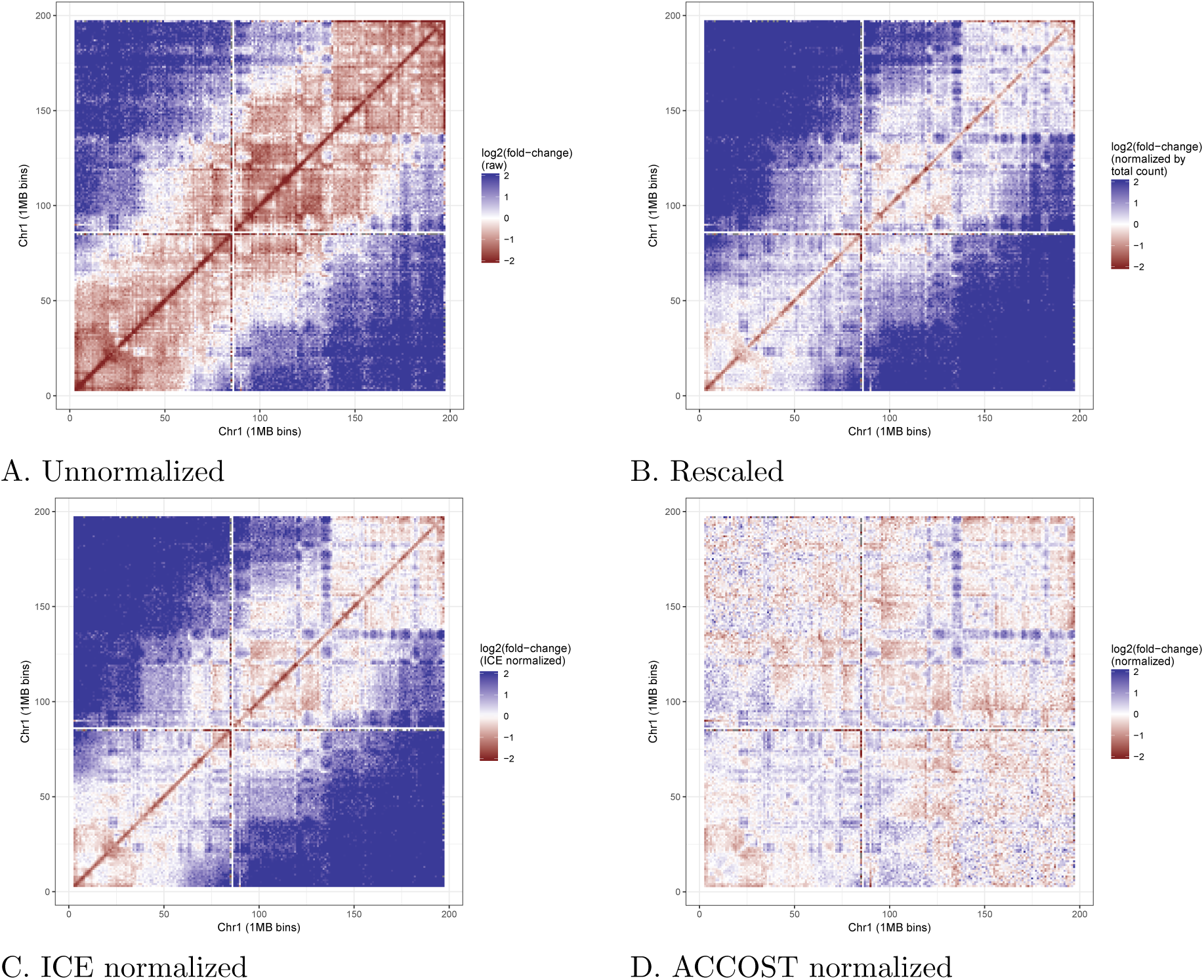
Genomic distance normalization highlights differences in contact counts. Each panel plots the log fold change of Hi-C contacts between ESC and cortex. (A) Raw data. (B) Data after rescaling the two matrices to have the same total count, to account for differences in sequencing depth. (C) Data after ICE normalization. (D) Data after ACCOST normalization, which additionally accounts for the genomic distance effect.

To quantitatively confirm that ACCOST correctly accounts for differences in the genomic distance effect, we simulated Hi-C matrices with different distance decay curves. We generated counts from Poisson distributions with means that decay as illustrated in Figure 3. Note that we did not employ a negative binomial distribution because we did not want the simulations to favor ACCOST. The significance of differences between simulated datasets was then assessed using ACCOST and three existing methods (diffHiC, FIND and HiCcompare). Because the only differences in the simulated data sets arise due to changes in the genomic distance effect, all methods should produce uniform (null) p-values. Empirically, we observe nearly uniform *p*-values for both ACCOST and diffHiC (Figure 4, top row). In contrast, both FIND and HiCcompare are misled by the genomic distance effect, with FIND assigning many very small *p*-values, and HiCCompare deviating both above and below the expected uniform distribution (Figure 4, bottom row).

**Figure 3:**
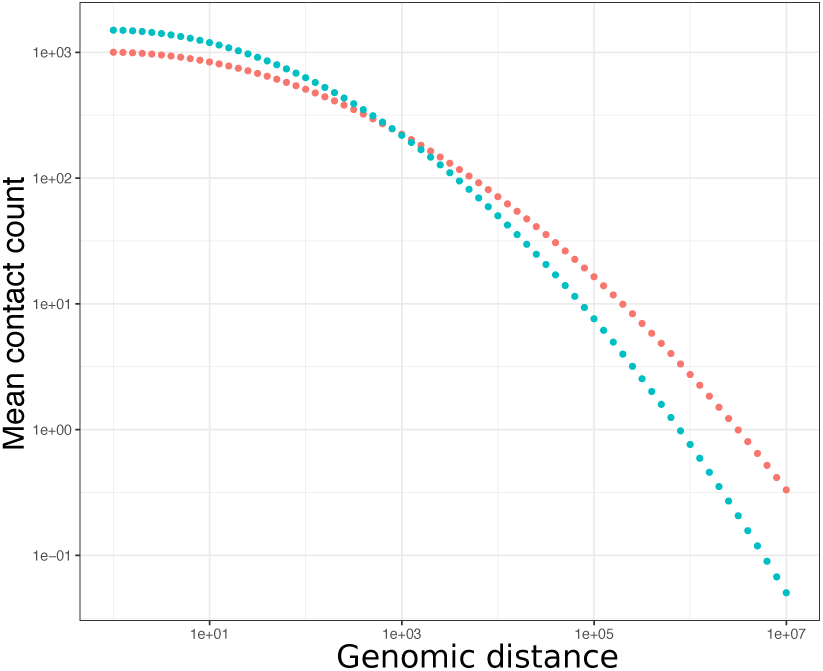
Simulated distance decay curves. The two decay curves were constructed such that the mean contact count between neighboring genomic bins was 1000 and 1500 for the two distributions. No position-specific biases (as are corrected for using ICE) were added.

**Figure 4:**
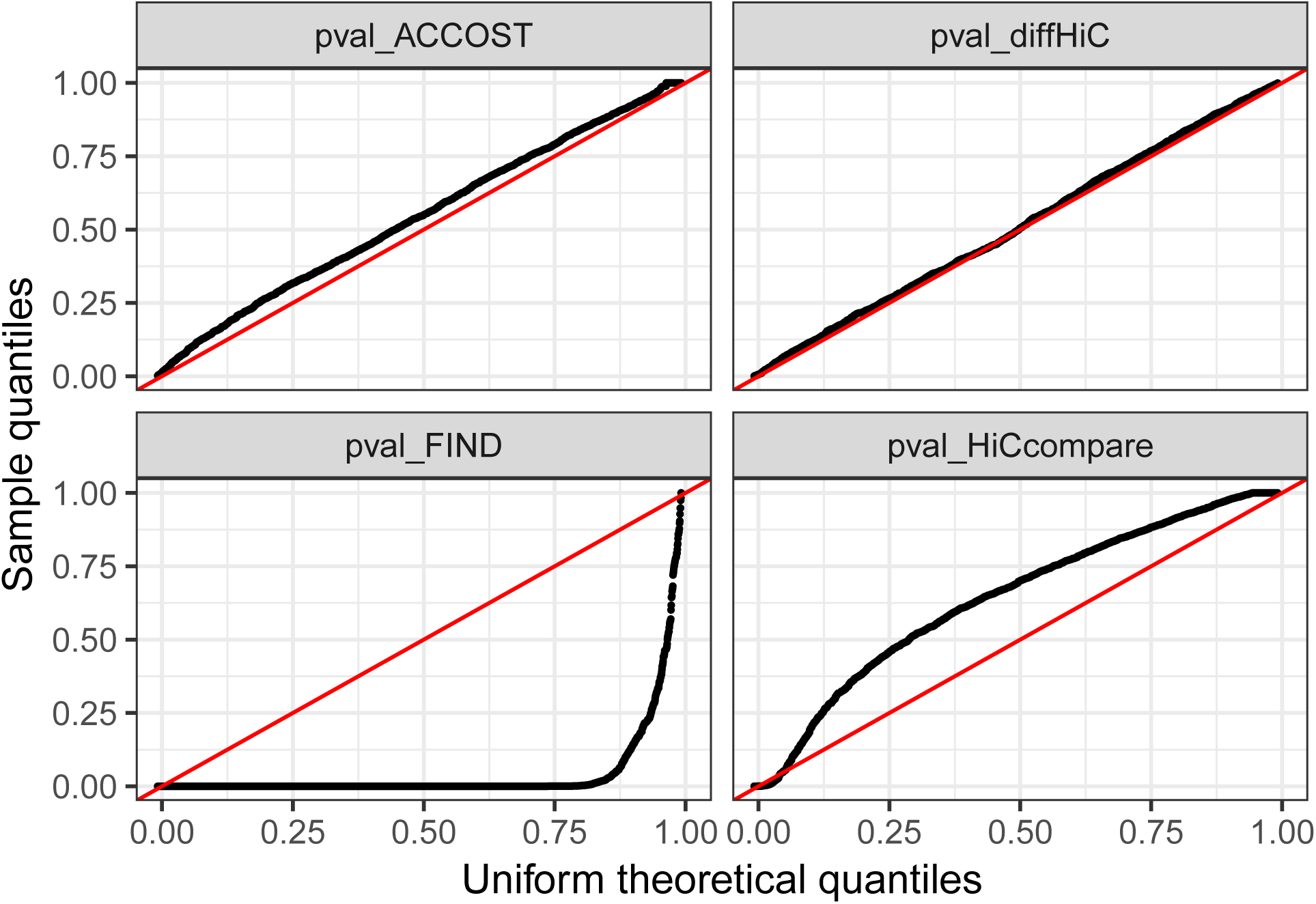
Uniformity of *p*-values generated by different methods. Each panel is a quantile-quantile (Q-Q) plot of the empirical (y-axis) versus uniform (x-axis) distribution of p-values generated by ACCOST, diffHiC, FIND, and HiCcompare on simulated null data. The *p*-values for FIND and HiCcompare deviate strongly from the expected uniform distribution.

We further compared ACCOST to diffHic by assessing the significance of changes in contact counts between GM12878 and IMR90 cells. These cells derive from two markedly different cell types (B-Lymphoeyte and fibroblast, respectively) and have been observed to have different short- and long-range contact patterns [19], consistent with differing genome organization at multiple scales. At short range (< 100 Mb, Fig. 5A and C), differential contact p-values are evenly distributed between positive and negative log fold-changes. However, at long range (> 100 Mb, Fig. 5B and D) diffHic p-values are skewed consistently toward more frequent contacts in IMR90 (negative log fold-change). Further investigation of this phenomenon shows a striking increase in the number of significant contacts called by diffHiC at genomic distance of 50 Mb (Fig. 5E), which corresponds to the threshold between the two LOESS normalizations performed by diffHiC.

**Figure 5:**
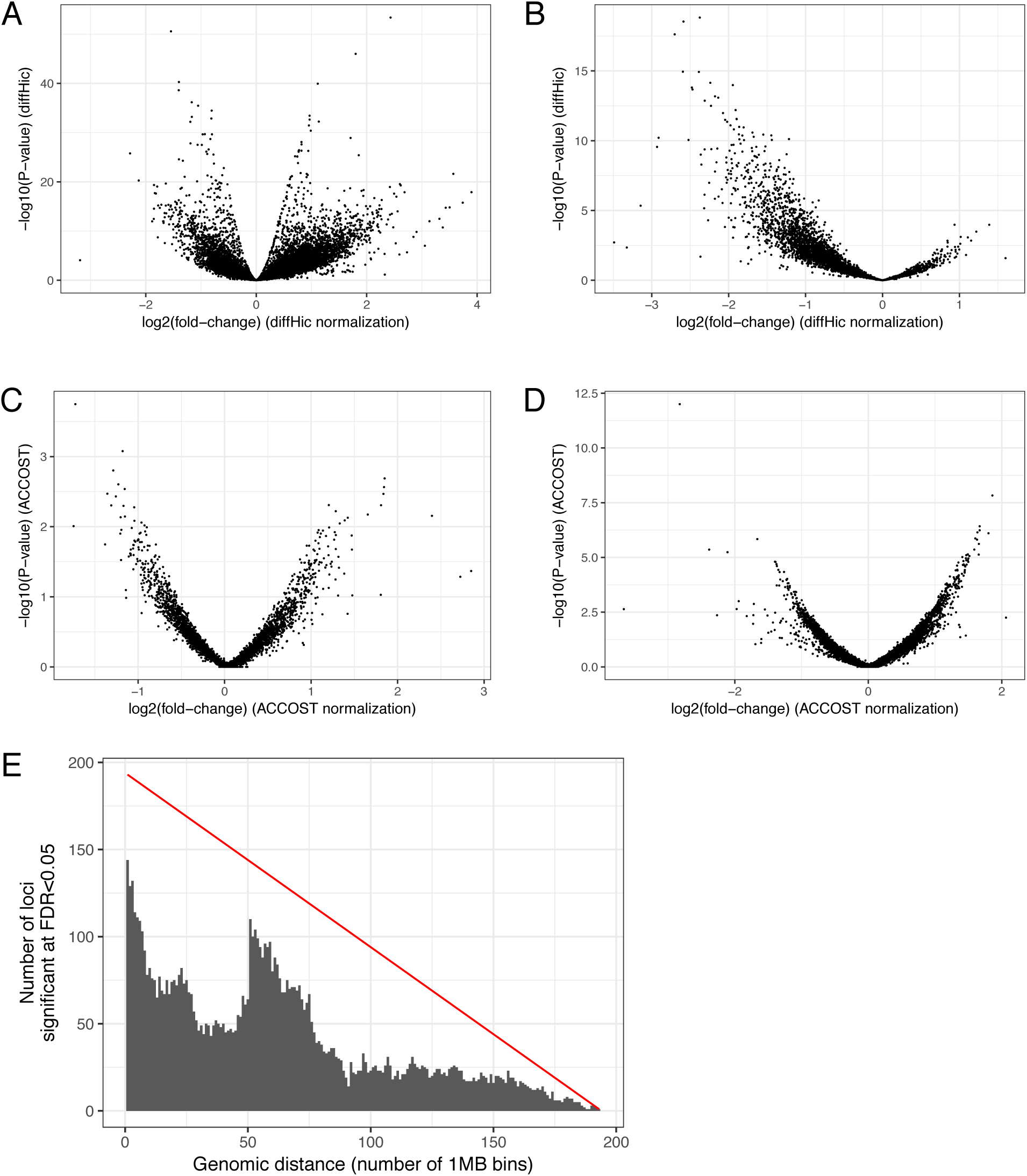
DiffHic p-values are skewed at long distances. (A and C) At short range (< 100 Mb), diffHic and ACCOST yield symmetric distributions of *p*-values. (B and D) At long range (> 100 Mb), diffHic fold-changes are skewed in one direction, while ACCOST fold-changes are more evenly distributed. (E) Histogram of bin pairs for which diffHic indicates a significant change in contact counts with FDR < 0.05. The red line indicates the total number of bin pairs measured.

### 3.2 Analysis of *Plasmodium* Hi-C data

As an illustration of how ACCOST might be applied in practice, we ran the software on Hi-C data from two stages in the *Plasmodium falciparum* life cycle: trophozoite, the most transcriptionally active stage in the erythrocytic cell cycle of the parasite, and sporozoite, the stage responsible for the transmission of the disease from mosquito to human. Malaria transmission specifically occurs when sporozoites are injected from the mosquito’s salivary gland into the human skin during blood feeding. ACCOST highlights several changes in interactions between these two stages, one of them particularly interesting in the right arm of chromosome 8 (Figure 6). In this chromosome, two clusters of genes interact significantly more at the sporozoite stage. One of these cluster involves genes specifically expressed at the sporozoite stage, including the sporozoite invasion-associated protein 2 and the sporozoite and liver stage tryptophan-rich protein. The generation of a loop at position ∼130k may be critical to enhancing gene expression of these genes at this particular stage. The other cluster of genes occurring a positions between 133k and 145k encode proteins that are exported to the surface of the infected red blood cells and implicated in mechanisms driving parasite immune evasion. These genes are critical to the parasite survival during the erythrocytic cycle and are expressed at a high level at the trophozoite stage. However, because these genes are not required in the mosquito stages, they are repressed in the sporozoite stage; accordingly, we observe them interacting significantly with the parasite heterochromatin cluster near the telomere ends at this stage of the parasite life cycle

**Figure 6:**
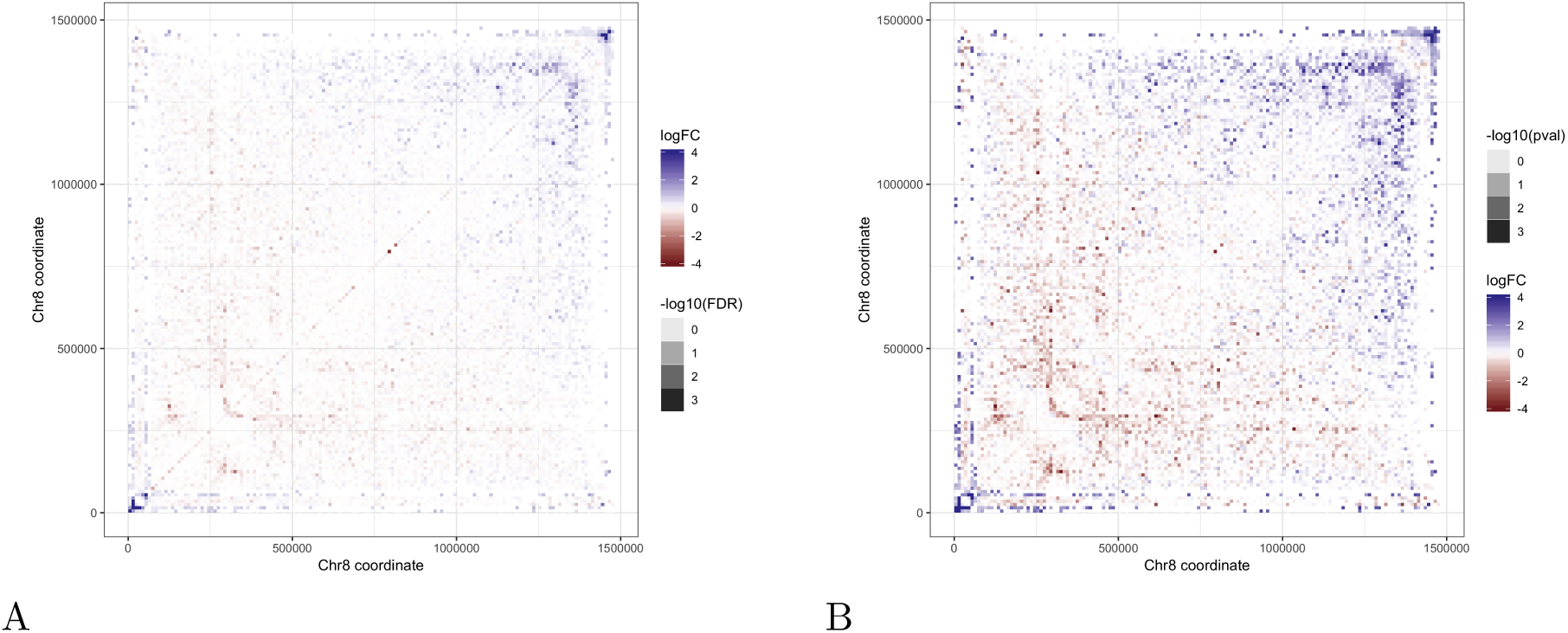
Comparison of life cycle stages in *Plasmodium falciparum.* (A) ICE-normalized contact counts from the end of chromosome 8 in the trophozoite stage (left) and sporozoite stage (right). (B) FDR-filtered fold changes highlight changes in chromatin conformation around clusters of genes (green bars) that are exported to the red blood cell surface during the trophozoite stage.

## 4 Discussion

We have demonstrated that ACCOST provides unbiased confidence estimates for differential Hi-C contacts by correctly controlling for the genomic distance effect caused by the polymer behavior of DNA. ACCOST thus enables biologists to better interpret Hi-C data by focusing on the subset of differential contacts that exhibit the most statistically significant differences. As is apparent from Figure 6, many of the observed differences exhibit complex structures related to known features of Hi-C data, such as loops, stripes, domains, and compartments. A clear direction for future work is to use ACCOST as a building block in a scheme to identify large, potentially irregularly shaped regions of differential contact. A second useful direction for future work would be extensions to handle differentiation or cell cycle time series data.

In the experiment reported in Section 3.1, the simulated counts follow a Poisson distribution, while much of ACCOST complexity is due to the fact that it assumes a negative binomial distribution. This mismatch is intentional, since we did not want our simulation setup to favor ACCOST, but the mismatch may also explain why the ACCOST p-values in Figure 4 are not perfectly uniform.

One surprising outcome of our simulation analysis is the conclusion that HiCcompare fails to acccount for the genomic distance effect, whereas diffHiC does a good job of handling the effect. We note that diffHic models the mean-variance relationship and then normalizes what it calls “trended biases” between libraries. Although the method never explicitly models the genomic distance, this trended bias seems empirically to capture the genomic distance effect. On the other hand, we do not have a good hypothesis for why HiCcompare does not work well. Similarly, the tendency of diffHiC to produce a greater number of significant p-values in one direction relative to the other remains unexplained (Figure 5).

As currently implemented, ACCOST uses one genomic distance bin per Hi-C bin. However, this is done merely for convenience. In principle, one could decouple these two discretization schemes. This would allow ACCOST, for example, to increase the size of genomic distance bins at long distances.

## Acknowledgments

This work was supported by NIH awards R01 AI136511 and U54 DK107979.

